# Cognitive Dynamics of Verb-Particle Constructions: An Eye-Tracking Study

**DOI:** 10.1101/2024.12.05.626940

**Authors:** Hassane Kissane, Konstantin Tziridis, Achim Schilling, Patrick Krauss, Thomas Herbst

## Abstract

This study investigates the cognitive processing of verb-particle constructions (VPCs) using eye-tracking data analysis method to explore how English native speakers process different types of the sequence NP–verb–particle–NP during reading tasks. While previous research has focused on phrasal verbs, our study extends this examination to include patterns with prepositions, aiming to identify distinct cognitive engagement patterns and processing efficiencies associated with each. We employed the Provo Corpus to analyse eye movements while participants read sentences containing these constructions. we focused on metrics such as first fixation duration, gaze duration, go-past times, and total reading times. Our findings indicate similarities in the lexical verbs, and significant differences in particles, indicating how these two types of constructions are processed, with phrasal verbs sometimes processed more efficiently than the prepositional counterparts. This suggests that phrasal verbs might be more deeply entrenched in the linguistic repertoire of native speakers, possibly functioning as single lexical units. This research contributes to the understanding of complex structures processing and the cognitive mechanisms that support it, offering insights that could influence linguistic theory and language education.

## 1. Introduction

### 1.1. The linguistic problem: identifying verb-particle combinations

Combinations of verb and particle have always attracted a lot of attention in the analysis and the teaching of the English language. One indication of this is the great number of so-called phrasal verb dictionaries flooding the market of English learners’ dictionaries – a type of dictionary that does not exist in this form for other Germanic languages. Since articles, adjectives and, with the exception of the genitive, nouns show no case marking in present-day English, the relations between the elements of clauses have to be expressed in other ways. Apart from word order, it is prepositions that play an important part in making syntactic relations clear. This is certainly one of the reasons why combinations of verbs with “small” function words such as *to, in* or *about* have attracted a lot of attention, another one being that many such combinations can be analyzed as being “idiomatic”, i.e., as expressing a meaning that cannot easily be derived from its component parts – just think of *look up* in the sense of consulting a dictionary or an encyclopaedia.

Despite (or because of) the importance of verb-particle combinations in present-day English, there is some terminological inconsistency in the use of the term *phrasal verb* and, indeed, there are various ways in the way such combinations have been analyzed. While phrasal verb dictionaries tend to have a very wide concept of phrasal verbs, others use the term only for one type of verb-particle combination. For instance, one of the standard reference grammars of the English language, the *Comprehensive Grammar of the English Language* by Quirk, Greenbaum, Leech, and Svartvik (1985) make a distinction between three main types, namely

i. prepositional verbs, in which the particle always precedes the noun phrase that these grammars as direct object as in:

(1)

a. I agree with Shawn. _TV-2010-Shawn_
b. agree Shawn with.
ii. phrasal verbs, in which the particle can precede or follow the noun phrase as in

(2)

a. … look up the word “semantics.” _TV-2010-Psych_
b. _b._ You can look the word up. _TV-2008-Psych_
iii. phrasal-prepositional verbs, which are combinations of a verb and two particles:

(3) I look forward to the opportunity to get to know you better. _TV-2012-Psych_

Not all linguistic models see things quite like this. The main difference between the approach taken by Quirk et al. (1985) and the valency model, which arose from Tesnière’s dependency theory and was applied widely in the description of German (e.g., Heringer 1996). In German, the concept of the prepositional object is one of the most common ways of analysing sequences, where valency theory has been extensively used more than in the analysis of English (Helbig 1991; Schumacher, 2004). However, it has recently been applied to the analysis of English (Herbst, Heath, Roe and Götz, 2004). This difference concerns the status of cases such as (1). Quirk et al. (1985) analyze (1) in terms of a prepositional verb which is followed by a direct object, whereas the valency approach sees the preposition as part of a prepositional phrase that functions as a complement of the verb (i.e., *agree + with Shawn.*) Identifying a category prepositional object enables us to put it in line with other complements, as in the case of *agree* (such as *agree* + *to do, agree* + *that-clause, agree* + *on-* phrase). Note that the other great reference grammar, the *Cambridge Grammar of the English Language*, edited by Huddleston and Pullum (2002: 274), makes use of the term prepositional verb for verbs which take prepositional complements without, however, regarding the preposition as a part of the verb.

What most, if not all, accounts of verb-particle constructions seem to agree on is that “accidental” co-occurrences of verbs with prepositions do not fall under the category of verb-particle combinations. In the examples under (1), the choice of preposition quite clearly is not determined by the verb – or, more technically, the valency properties (Helbig & Schenkel 1973; Herbst et al. 2004; Herbst & Schüller 2008) or the subcategorization frame of the verb (Chomsky 1965: 190) or the argument structure constructions the verb participates in (Goldberg 2019). Rather, the prepositional phrase is part of an adjunct construction which does not form a unit with the verb in question, but can appear with more or less any verb (as long as it makes sense). Typical examples are locative and temporal adjunct constructions, i.e., expressions indicating the location of someone or something in space. Compare:

(4)

a. They met/married/honeymooned/stayed/lived in New York.
b. They met in Sweden/at the White House/outside the courtroom/under the bridge/on a tram.

Basically, what all of these different classifications do is to capture different degrees of association between a verb and a particle. Leaving aside intransitive phrasal verbs (***The plane took off***) and combinations with two particles for the moment, we can distinguish between the following (prototypical) cases:

**Table 1.** Types of verb + particle + noun phrase combinations in terms of structure (NP: noun phrase; PP = prepositional phrase, i.e. particle + noun phrase)

|  |  | The particle is |  |  |
| --- | --- | --- | --- | --- |
| A | verb + PP-adjunct | not determined by verb<br>valency | not<br>shiftable | not<br>stressed |
| B | verb + PP-object | determined by verb valency | not<br>shiftable | unstressed |
| C | verb + part +<br>NP | determined by verb valency | shiftable | stressed |

From a purely **structural point of view**, it would seem that the association between a verb and a particle is weakest in the case of A, especially since PP-adjuncts can also occur clause initially, as in (5):

(5) In the United States, only 1 percent of trips are made by bicycle. In the Netherlands, which has only 1/18 of the U.S.’s population, that number is close to 26 percent._COCA-2013-MAG_

From a **usage-based point of view**, however, we must also allow for other factors influencing the strength of association such as frequency and meaning. In the *Corpus of Contemporary English* (COCA), the verb form ***lived*** is followed by the word form ***in*** in over 30% of all cases (based on COCA: ACAD, FIC, MAG, NEWS, SPOK). The transitional probabilities (Jurafsky, 2003) do not necessarily coincide with the degree of affinity one might expect on the basis of the structural classification:

**Table 2.** Transitional probabilities of the sequences *agree on, look up* and *live with* according to the Corpus of Contemporary English (subcorpora ACAD, FIC, MAG, NEWS, SPOK)

|  | agree on | look up | live in |
| --- | --- | --- | --- |
| verb + part | 6773 | 29356 | 83293 |
| verb total | 114481 | 844409 | 293032 |
| transitional probability | 0,05916266 | 0,03476514 | 0,28424541 |

When determining the lexical status of verb-particle combinations, i.e. the question of whether it is appropriate to regard a particular verb-particle combination as a unit in the lexicon, i.e. a lexeme in the sense of Cruse (1986:80), the factor of meaning plays an essential role. However, as with transitional probability, there is no 1:1-correspondence of the degree of idiomaticity and the three formal criteria described above: The examples under (6) and (7) are both idiomatic in the sense that their meanings cannot be deduced easily from their component parts, but the particle can only be shifted in the case of (7):

(6) She’s to look after you in case anything should happen to the two of us. _TV-2007-Psych_
(7)

a. But why in hell should the police make up a story about an accident? _BNC-A7A-1487_
b. Why would he make the story up? _BNC-CBF-173_

Furthermore, it has long been recognized that there are various degrees of idiomaticity in such cases; see, e.g. Herbst’s (2010: 135–136) discussion of examples such as those under (8) in terms of a “scale of idiomaticity” (see also Cowie & Mackin 1975: ix-xii):

(8)

a. …when Marge Simpson put the cat out. _COCA-1994-TV_ [neither *put* nor *out* used idiomatically]]
b. Eat it all up. It will make you strong. _TV-1966-Jeanniie_ [*up* used in the sense of *end-point*]
c. Shawn, you made that name up one minute ago. _TV-2010-Psych_ [*up* used idiosyncratically]
d. But he insisted that I give up smoking … _COCA-2017-NEWS_ [neither verb nor particle used idiomatically]

A strong degree of semantic non-transparency is a convincing argument for attributing lexeme-status to a combination. Since syntactic idiosyncrasies (such as the fact that ***the bucket was kicked*** can only be used in a jocular way), it is not surprising that phrasal verbs were treated as a special type of verbs in many approaches (Fraser, 1965/1966/1976; Görlach, 2004), also, Lipka (1972), who stated that this category of verbs “can be regarded as a particular surface structure shared by a large number of lexical items with various word-formative and semantic structures.” (13-17). However, it must be underscored that this does not mean that phrasal verbs are a clearly defined category in that, as the lexicographical description provided by Cowie & Mackin (1975) in the *Oxford Dictionary of Current Idiomatic English* (ODCIE) has shown, by no means all members of the category share all of its features.

As mentioned above, verb-particle combinations span a range from transparent to opaque meanings. Some phrasal verbs, such as ***put out***, can be deciphered from the literal meanings of their components. In contrast, others like ***break up*** require an idiomatic understanding. Such idiomatic interpretations, anchored in linguistic contexts, often elude direct translations into other languages. A single-word synonym could replace some, however. As proposed in the ODCIE, to ascertain whether a phrasal verb is idiomatic, the phrasal verbs construction should be:

- Replaceable by other single words with similar meanings, e.g., ***Bill had to put off the meeting until next week***, the phrasal verb ***put off*** can be replaced with ***delay*** or ***postpone***.
- If the phrasal verb is idiomatic, its components should be irreplaceable with words of like meaning, e.g.:

(9)

a. John took off Winston Churchill to perfection. Here, the verb “took” cannot be replaced with similar-meaning verbs while retaining the same overall meaning.
b. *John snatched off Winston Churchill to perfection.
- The convertibility into nouns, for instance, the phrasal verbs ***break down*** and ***make up*** have corresponding nouns ***breakdown*** and ***make-up***.

The first measure of idiomaticity for phrasal verbs can also be applied to idiomatic verb-preposition combinations such as ***take after***. The entire combination can be substituted with the verb ***resemble*** based on the first criterion. Moreover, the main verb cannot be replaced with other semantically similar verbs:

(10)

a. I took after my mother. We are both impatient.
b. I resemble my mother. We are both impatient.
c. *I snatched after my mother. We are both impatient.

The key issue that we wish to address here is whether there is any cognitive and neurolinguistic evidence to support the claim that what Quirk et al. (1985) call prepositional verbs are units of the lexicon in the same way that phrasal verbs are or to support the view propagated by valency theory and also Huddleston and Pullum (2002) that prepositional phrases are complements of the verb in very much the same way that noun phrases, *that*-clauses or *to-*infinitives are.

### 1.2. Verb-Particle Combinations in Eye-tracking Studies

Eye-tracking studies directed to verb-particle combinations have emerged as a research method in both linguistics and psycholinguistics. These studies, by examining the processing and comprehension of verb-particle combinations, lighten the cognitive processes regarding language production and comprehension. Herbay et al. (2018) examined how French-English bilinguals process Verb-Particle Constructions (VPCs) within sentences, drawing on a framework proposed by Gonnerman and Hayes (2005). They investigated whether bilingual processing parallels that of monolinguals, emphasizing the particle position, NP length, and VPC dependency. Their findings clarified the influence of VPC lexical knowledge and working memory on reading patterns. Bilinguals with limited VPC comprehension showed unexperienced processing, whereas those more proficient in VPC displayed native-like reading behaviors. This study investigated the significance of VPC lexical knowledge in sentence processing since deficits in this knowledge can block syntactic processing in second-language sentence understanding.

In another study, Tiv et al. (2019) explored the comprehension of VPC among English-French bilingual adults, in which they considered these combinations as a domain recognized for its inherent challenges for second-language (L2) learners. The results showed that native (L1) readers typically access lexicalized VPC representations directly. Particularly, monolingual readers often find the adjacent forms more intuitive, and they process semantically transparent VPCs more rapidly than the opaque ones. it has been observed that the use frequency of a particular VPC might determine its reading pace. Moreover, L1 English readers did not present a clear preference for either the adjacent or split particle positions. This observation indicated that other item-specific attributes, such as frequency and co-occurrence strength, possibly modulate reading patterns. In contrast, L2 readers engaged in more compositional processing. Therefore, several factors have shaped the VPC processing, including the form (adjacent or split), the degree of semantic transparency, and frequency.

While prior eye-tracking research on verb particle combinations (VPCs) in English monolinguals and bilinguals by Tiv et al. (2019) and Herbay et al. (2018) focused on phrasal verbs, there was an absence of attention to verb-preposition combinations. Recognizing this gap, our study aims to comparatively assess the cognitive processing associated with both phrasal verbs and verb-preposition combinations. the study objective is to identify how linguistic nuances between these two subtypes influence cognitive engagement during reading, aiming for a comprehensive understanding of VPCs.

Using eye-tracking, we examine eye movements as participants processed these constructions, investigating real-time cognitive interactions. Our focus extended to the role of frequency, length, and predictability in VPC processing, seeking to identify if these parameters modulate the ease with which one form is processed over the other. If this holds true, some VPCs, conditioned by these factors, might be deeply entrenched in the reader’s linguistic repertoire, functioning more as singular terms rather than compound structures. Our overarching aim is to demystify the cognitive tactics English readers employ when navigating verb-particle constructions.

The core research queries we sought to address are:

a. How do native English speakers’ eye movements differ when processing phrasal verbs as opposed to verb-preposition combinations during reading tasks? Is there a processing advantage for phrasal verbs over verb-preposition combinations? Should phrasal verbs alone be perceived as single lexical entities, efficient processing should be more evident in these compared to verb-preposition combinations.
b. How do the length, frequency, and predictability of VPCs collectively impact both early and late eye movement measures during the reading of sentences featuring VPCs?

## 2. Methods

### 2.1 Original dataset (Provo corpus)

In this study, we use Provo eye-tracking Corpus (Luke & Christianson, 2018). The Provo Corpus consist of Fifty-five short passages that were sourced from various materials like news articles, science magazines and fiction, averaging 50 words in length (range: 39–62) with 2.5 sentences each (range: 1–5). Each sentence had about 13.3 words (range: 3–52). In total, the passages had 2,689 words, with 1,197 unique word forms. The eye-tracking data that this study used was extracted from the previously mentioned corpus, which was collected from an experiment that recruited native American English speakers (n=84), who had 20/20 corrected or uncorrected vision. Participants were presented with the written passages. The passages were presented one at a time on a computer screen, with eye-tracking technology used to measure the participants’ eye movements when they read the sentences.

The data were collected with an SR Research EyeLink 1000 plus eye-tracker sampled at 1000 Hz (for the full description of the method and stimuli, see Luke & Christianson, 2017). This corpus was ideal for our purpose as it contained a reasonable number of verb-particle combinations with variant categories, and the eye movement measures were provided for all the words in the corpus.

### 2. 2. Extracted regions of Interest

In the context of our investigation, we directed our focus towards phrasal verbs and verb-preposition combinations, which in regular language use are often used in more complex syntactic structures. To make the target words analysis simple, we extracted these specific types of verb-particle constructions from the larger corpus. This enabled the analysis to concentrate on the fundamental lexical verbs and the particles involved in these constructions’ formation. The extracted dataset Consists of a total of 19 VPCs, which were further categorized into two subsets: 10 phrasal verbs 9 verb-preposition combinations. The phrasal verbs in the corpus were comprised of lexical elements tagged as “Verb” paired with adverbial particles tagged as “Adverb”. However, an tagging mistake was observed, wherein two particles were classified as “Preposition” instead of their accurate “Adverb” categorization. In parallel, the verb-preposition combinations included in our study were characterized by lexical verbs tagged as “Verb” which were conjoined with prepositional particles annotated as “Preposition”. The VPCs also ranged on other aspects (e.g., verb tenses) that was not directly relevant to the current study.

### 2.3. Eye Tracking Measures

Eye tracking systems record several measures that represent the eye reactions when dealing with a specific word or region of interest. These measures are separated into two groups, early and late measures. Early measures of reading are posited to capture preliminary, pre-semantic stages of lexical access within memory, specifically focusing on the retrieval of a word’s form devoid of semantic representation. In contrast, late measures are theorized to reflect subsequent semantic processes, including the understanding of a lexical item and its contextual integration within the context. (Haeuser, Baum, & Titone, 2021).

In this work, we focus on three early measures: First fixation duration (The duration of the first fixation on the interest area, in milliseconds), which primarily stands for lexical information processing, like lexical access (Inhoff, 1984) and is usually affected by lexical factors such as length, frequency and predictability (Staub and Rayner 2007; Carrol and Conklin, 2015). Gaze duration / first pass reading time (the sum of the duration across all fixations) of the first run within a lexical verb, which is a measure for lexical access, that computes the sum of the duration of individual fixations before moving to the next region of interest (Inhoff and Radach, 1998; Rayner, 1998; Salicchi, Chersoni, & Lenci, 2023). Go-past times (the summed fixation duration from when the reader looks at a target word as well as any time spent rereading earlier words in the sentence before moving to the next words) of the particle whether adverbial or prepositional (Haeuser, Baum, & Titone, 2021). Late measures: total reading time (i.e., summation of the duration across all fixations) on both the lexical verb and the following particle separately. This measure reflects more strategic, controlled processes involved in reading comprehension and is affected by both efficiency of lexical access and sentence-level processing, indicating a high semantic representation of the word within sentence context (Radach & Kennedy, 2013; Salicchi, Chersoni, & Lenci, 2023).

### 2.4. Length, Frequency, and Predictability Testing

While there are various English Corpora for linguistic pattern research, we used the Corpus of Contemporary American English (COCA) for the frequency analysis considering that the eye-tracking data is collected from native American English speakers. Thus, other corpora like the British national corpus would not be the perfect resource to define the correlations between the word frequencies and eye-tracking measures. The analysis investigated the frequency of verb, verb-particle combinations, and individual particles.

Regarding predictability, there is a growing interest in using the language models’ capabilities to assess the linguistic predictabilities through tasks like *next-word prediction* with recurrent neural networks such as LSTM (Long Short-Term Memory) (Hochreiter and Schmidhuber, 1997), and *fill-in the blank task* with pre-trained language models such as BERT (Bidirectional Encoder Representations from Transformers) (Devlin et al., 2019). In our test, we use the pre-trained transformers-based language model BERT to investigate the prediction probabilities of verbs in natural language sentences. We used the original sentences that were showed to the participants for the eye-tracking data acquisition and systematically masked the target verbs. Therefore, we presented these modified sentences to BERT and tasked it with predicting the masked verbs with their prediction probabilities.

**Figure 1:**
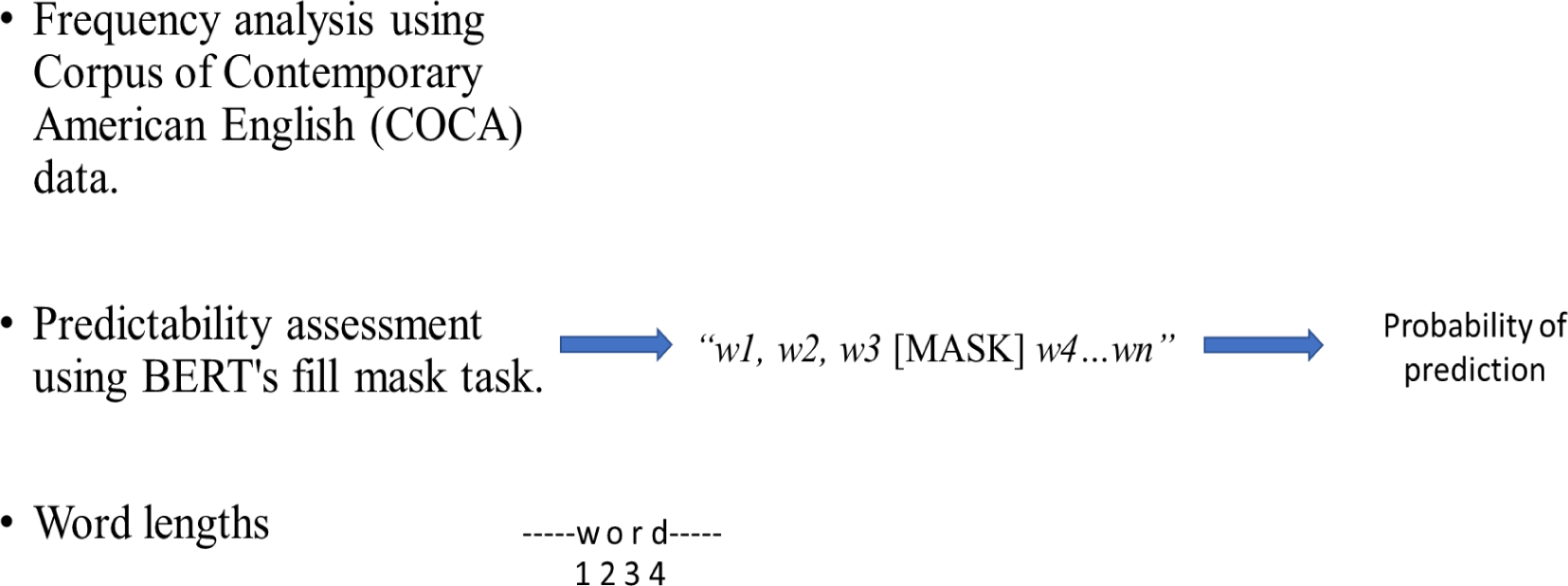
The procedures to define Frequency, predictabilities and lengths of VPCs as usage-based information for analysed constructions.

### 2.5. Statistical testing

For our statistical analyses, we first tested the normal distribution of data for both the eye-tracking measures and usage-based factors using the Kolmogorov-Smirnov test, which reported significant deviations from normality across all assessed measures and factors. The statistics and corresponding p-values indicated substantial non-normality (see Table 3). Thus, we used a non-parametric method for hypothesis testing. We computed separate Mann-Whitney U tests (Mann & Whitney 1947; McKnight & Najab, 2010) for both the eye-tracking measures and usage-based factors; each variable is being computed with respect to the category of the VPCs (PV: phrasal verb and PRP: verb-preposition combination). Therefore, we run a non-parametric correlation analysis with Spearman correlation to detect any potential correlation between the eye movement patterns and the usage-based information of the defined VPC. All the statistics done with DATAtab (DATAtab Team, 2023), provided online through a paid subscription.

**Table3:** Kolmogorov-Smirnov test for normal distribution of data in eye tracking measures and usage-based factors.

| Measure/Factor |  | Statistics | p |
| --- | --- | --- | --- |
| Eye tracking measures | First fixation duration | 0.11 | <.001 |
|  | Gaze duration | 0.14 | <.001 |
|  | Go-past times | 0.28 | <.001 |
|  | Total reading time (verbs) | 0.17 | <.001 |
|  | Total reading time (particles) | 0.19 | <.001 |
| Usage-based factors | Length | 0.36 | <.001 |
|  | Frequency | 0.22 | <.001 |
|  | Predictability | 0.21 | <.001 |

## 3. Results

### 3.1. Eye-tracking results

The results showed that the reading times, as represented in the first fixation duration, did not differ between the two verb types; the median of the first fixation duration for both categories were very similar, with 200 ms for phrasal verbs and 194 ms for verb-preposition combinations. The Mann-Whitney U test (U = 173357.5; p = 0.543) reported no difference between the two categories with respect to the dependent variable, first fixation duration.

Similarly, another metric, gaze duration, did not show differences; participants spent approximately the same amount of time on words from phrasal verbs as they did on words from verb-preposition combinations. The median gaze duration for phrasal verbs was 208 ms, while for verb-preposition combinations it was 207 ms. the statistical test reported (U =171549; p = 0.361), indicating that there is no statistically significant difference between the two categories in respect to the gaze duration variable.

The median go-past times for particles were different; 191 ms for phrasal verbs and 200 ms for verb-preposition combinations. Nevertheless, they did not show a statistically significant difference (U = 75445; p = 0.199).

**Figure 2:**
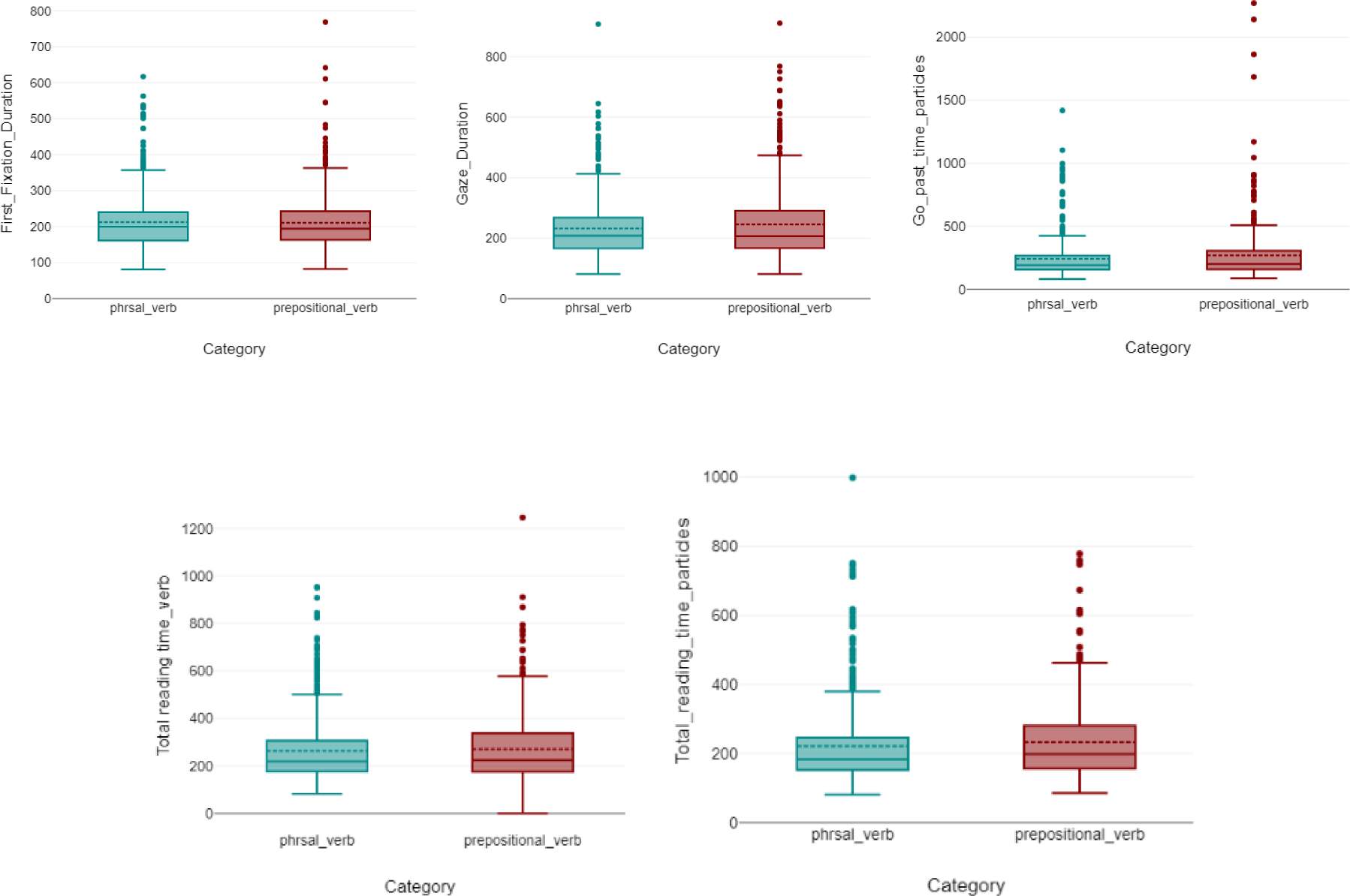
Visualizations of first fixation duration, gaze duration, go-past times, and total reading time of lexical verbs and particles for phrasal verbs versus verb-preposition combinations, showing similar median values and no significant differences in the metrics except for total reading times in particles.

Consequently, the late measure standing for semantic representation and the word understanding within the sentence which is total reading time showed no significant differences; Participants reported similar reading times while dealing with the distinct categories. The median reading time for lexical verbs in phrasal verbs was 219 ms while for verb-preposition combinations was 223.5 ms. Therefore, this measure showed non statistically significant difference (U = 171765; p = 0.354) regarding the lexical verbs.

However, the total reading times of particles showed significant differences; Participants reported efficiency reading phrasal verbs in times less than verb-preposition combinations. The median for particles in phrasal verbs was 184 ms while for verb-preposition combinations was 198.5 ms. Therefore, the total reading time measure showed a statistically significant difference between the particles in the two distinct categories (U = 72563.5; p = 0.023).

### 3.2. Length, frequency and predictability as usage-based information for the defined VPC

These results, while indicating some notable cognitive and linguistic similarities in the processing of phrasal verbs and verb-preposition combinations, may also be influenced by external factors, such as the length, frequency and predictability of the verb constructions. This is what the second question addresses. Therefore, we test another hypothesis on the equality of the two groups in terms of these external factors.

The statistical analysis indicated a statistically significant difference between the two types of constructions (U = 218736; p < 0.001), suggesting that there is a significant variation in the length of phrasal verbs compared to verb-preposition combinations. The results of the predictability test also showed a statistically significant difference between the two verb types (U = 232848; p < 0.001), indicating that there is a discernible variation in predictability between phrasal and verb-preposition combinations. However, the frequency of the two categories of these constructions did not demonstrate a significant difference, as evidenced by (*U* = 317520; *p* = 1).

**Figure 3:**
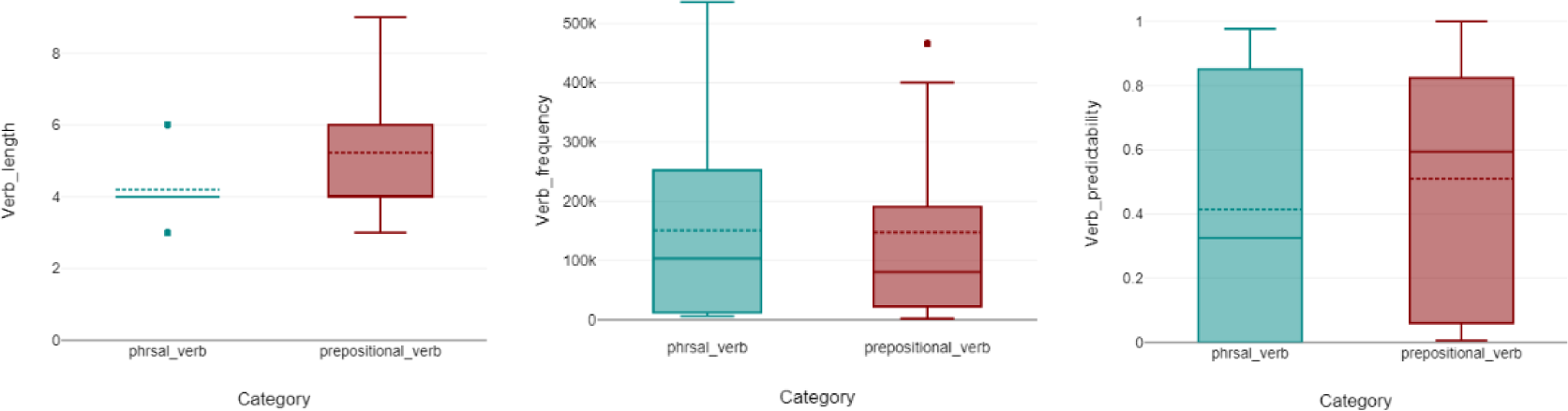
visualizations of the usage-based information; length frequency and predictability.

The most remarkable correlation between the usage-based information and eye movement measures is observed in the length of the verb construction with total reading time, where the highest significant positive correlation has been reported in phrasal verbs with an (*r = 0.21, p = 0.001*). A negative correlation is noted in phrasal verbs between the higher predictability that correlates with the short total reading times by (*r = −0.19, p< 0.001*). However, there was no correlation between the verb in verb-preposition combinations category and this eye movement measure.

**Table 4:**
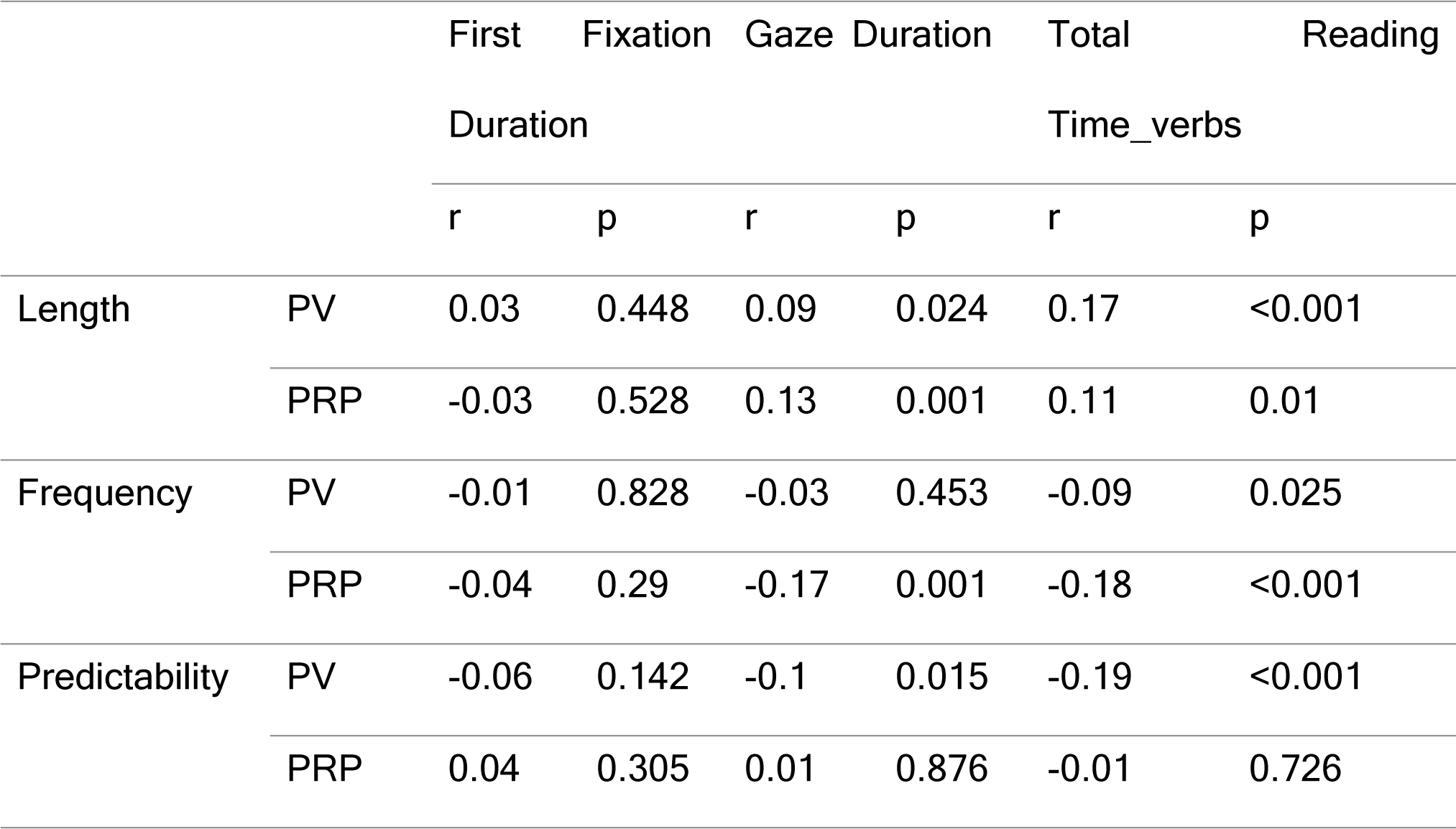
Spearman Correlation Analysis.

## 4. Discussion

This study aimed to clarify the language processing mechanisms during continuous speech comprehension by measuring behavioural data (eye movements) while conducting continuous sentence reading. By determining the category of the constructions, we discussed the linguistic and cognitive mechanisms in processing sequences of verb + particle + NP within a valency framework. In this study, the investigation of the interaction between linguistic features and eye-tracking measures has shown the cognitive processes underlying the reading of phrasal verbs and verb-preposition combinations. Specifically, the behavioural interactions of native speakers to these verbs through their eye movements provide an understanding of the dynamics of reading. Lastly, we examined whether these eye movement patterns were influenced by usage-based factors; length, frequency, and predictability, and how they illustrate these linguistic properties and their interaction with eye movement patterns. Therefore, we evaluated the accuracy of several machine learning models on the classification task of those verb patterns based on the acquired eye movement data while reading them.

### Phrasal verbs versus verb-preposition combinations

#### Predictability and efficient lexical access in phrasal verbs

The results of the study demonstrate a negative correlation between predictability and eye-tracking measures for phrasal verbs. Specifically, predictable phrasal verbs are associated with shorter total reading times. This pattern suggests that, in the context of our experiment, phrasal verbs with higher predictability are accessed and processed more efficiently. Therefore, this finding might align with linguistic claims proposing that phrasal verbs are single lexical units (Cappelle, Shtyrov & Pulvermüller, 2010; Pulvermüller, Cappelle & Shtyrov, 2013: 415).

#### No Clear Predictability Effects in verb-preposition combinations

Contrastingly, for verb-preposition combinations, the correlations between predictability and eye-tracking measures are not statistically significant. This lack of a clear relationship indicates that predictability may not strongly influence visual processing efficiency for this specific linguistic category in our study. The absence of a predictive effect challenges the notion that predictability uniformly shapes processing across different types of verbs, highlighting the nuanced nature of linguistic processing. Therefore, what is very important with these results is that the verbs are more predictable in verb-preposition combinations than in phrasal verbs, but predictability is only correlated with phrasal verbs but not with verb-preposition combinations.

#### Connection with past research

In the context of the neurolinguistic experiments that suggest phrasal verbs are processed as single lexical units in the brain (Cappelle, Shtyrov & Pulvermüller, 2010; Pulvermüller, Cappelle & Shtyrov, 2013: 415), our study aimed to contribute to the ongoing discourse surrounding the processing of verb-particle combinations. Traditional grammar models (Quirk et al,1985) often treat both phrasal and verb-preposition combinations as single lexical units, exemplified in sentences like ***She looked after her son,*** where the sequence ***looked after*** is considered a single lexical unit called ***prepositional verb***. However, the valency theory interpretation, proposed by Herbst and Schüller (2008), introduces an alternative perspective on analysing these combinations. In this approach, valency relates to the number and types of arguments that a predicate, mainly a verb, can govern. Considering verb-preposition combinations like ***decide on***, ***decide against***, and ***decide in favour of***, the valency theory provides an alternative interpretation; instead of viewing these combinations as single lexical units, as some grammatical frameworks might (Quirk et al,1985), valency theory perceives the main verb ***decide*** and its prepositional complements (such as ***on****, **against**, **in favour of**)* as separate yet fundamentally interconnected elements that contribute to the predicate’s valency. For instance, in the sentence:

##### - They decided on the bus

The verb ***decide*** is considered as the predicator. Meanwhile, the prepositional noun phrase ***on the bus*** is identified as one of the potential complement classes fulfilling the verb’s valency requirements. This interpretation offers multiple benefits, such as consistent formalization across various usages of the verb and its complements. Crucially, it avoids unnecessary expansion of the number of lexical units in consideration, enabling a systematic description based on the semantic roles of different complement classes. This perspective supports the comprehension of the intricate syntactic relationships between the verb, its prepositional complement, and other sentence components.

In our study and the valency theory framework context, our findings did not strongly support a distinct processing pattern between phrasal verbs and verb-preposition combinations regarding early eye-tracking measures. However, a difference in a late measure (total reading times of particles) is provided. While neurolinguistic experiments have suggested a unique neural processing mechanism for phrasal verbs, our behavioural measurement did only differentiate the processing of phrasal verbs and verb-preposition combinations at the higher level of particles processing which is usually connected with semantic level processing. Valency theory, which posits a more segmented processing approach for verb-preposition combinations, did partially find empirical support in our study.

These results did not fully support the strict application of the valency theory to the cognitive processing of phrasal verbs and verb-preposition combinations during reading through the early measures in both lexical verbs and particles and the late measures in lexical verbs. However, in late measures regarding particles, there was a significant difference between the two categories, with reported longer reading times on particles in verb-preposition combinations than in phrasal verbs. These results would interact with the “word group” hypothesis, which argues that multiple words elicit eye-movement patterns that are typical for single words (Radach, 1996; Kliegl, Risse, & Laubrock, 2007). This insight from the ‘word group’ hypothesis could be used to explain why the particle total reading times are shorter in phrasal verbs than in verb-preposition combinations, suggesting that this is because they are involved in single lexical units with the main verb. Therefore, they receive less attention because the grouping of the lexical verb and the following particle results in a reduction in the time spent reading this type of linguistic combinations.

In conclusion, our study contributes to the broader discussion on the processing of verb-particle combinations by highlighting the challenges in reconciling theoretical linguistic frameworks, such as valency theory, with empirical behavioural measures. Further research is warranted to explore the interplay between theoretical linguistics and cognitive processing, considering additional linguistic features and experimental conditions that may influence the observed patterns.

In addition to exploring the influence of proficiency on phrasal verb processing mechanisms, future studies could also investigate the role of the semantic states of these constructions (literal, semi-idiomatic, and fully idiomatic). In the present investigation, phrasal verbs and verb-preposition combinations were embedded in natural sentences of continuous speech; however, one might ask whether the apparent preference for figurative interpretations is upheld when introducing biased contextual information. In other words, will the figurative meaning of phrasal verbs be accessed when the sentence or discourse context is biased towards the literal interpretation?

## 5. Acknowledgements

This work was funded by the Deutsche Forschungsgemeinschaft (DFG, German Research Foundation): grants KR 5148/5-1 (project number 542747151), KR 5148/3-1 (project number 510395418) and GRK 2839 (project number 468527017) to PK and TH, grant TZ 100/2-1 (project number 510395418) to KT, and grant SCHI 1482/3-1 (project number 451810794) to AS.

## 6. Data Availability Statement

In this study, we used an open-access data set, the Provo eye-tracking Corpus (Luke & Christianson, 2018), which is freely available.

## 7. Ethical Approval

No ethical approval was necessary for this study, since we used an already existing open-access data set and did not perform any new measurements or experiments with human.

## Notes

### Competing Interest Statement

The authors have declared no competing interest.

